# Differential analysis of pesticides biodegradation in soil using conventional and high-throughput technology

**DOI:** 10.1101/2021.06.01.446544

**Authors:** Saurabh Gangola, Samiksha Joshi, Saurabh Kumar, Anita Sharma

## Abstract

A potential pesticide degrading bacterial isolate (2D), showing maximum tolerance (450 ppm) for cypermethrin, fipronil, imidacloprid and sulfosulfuron was recovered from a pesticide contaminated agricultural field. The isolate degraded cypermethrin, imidacloprid, fipronil and sulfosulfuron in minimal salt medium with 94, 91, 89 and 86 % respectively as revealed by HPLC and GC analysis after 15 days of incubation. Presence of cyclobutane, pyrrolidine, chloroacetic acid, formic acid and decyl ester as major intermediate metabolites of cypermethrin biodegradation was observed in GC-MS analysis. Results based on 16S rDNA sequencing, and phylogenetic analysis showed maximum similarity of 2D with *Bacillus cereus* (MH341691). Stress responsive and catabolic/ pesticide degrading proteins were over expressed in the presence of cypermethrin in bacteria. Enzyme kinetics of laccase was deduced in the test isolate under normal and pesticide stress conditions. Amplification of laccase gene showed a major band of 1200bp. Maximum copy number of 16S rDNA was seenin uncontaminated soil as compared to pesticide contaminated soil using qRT-PCR. The metagenome sequencing revealed reduction in the population of proteobacteria in contaminated soil as compared to uncontaminated soil but showed dominance of actinobacteria, firmicutes and bacteriodates in pesticide spiked soil. Presence of some new phyla like chloroflexi, planctomycetes, verrucomicrobia was observed followed by extinction of acidobacteria and crenarchaeota in spiked soil. The present study highlights on the potential of 2D bacterial strain i.e., high tolerance level of pesticide, effective biodegradation rate, and presence of laccase gene in bacterial strain 2D, could become a potential biological agent for large-scale treatment of mixture of pesticide (cypermethrin, fipronil, imidacloprid and sulfosulfuron) in natural environment (soil and water).

## Introduction

From the last 20th century the overall global production of grain has increased from 500 million tons to 700 million tons [1]. However India alone produces 250 million tons of grain but it also losses 11 to 15% of total produced grains per year due to the pests attack and by some other causes [2]. To prevent such losses and to achieve the food targets, farmers have adopted to use hybrid seeds, systematic irrigation and application of chemical fertilizers and pesticides [3–4]. Cypermethrin, fipronil, sulfosulfuron, imidacloprid are some common pesticides used widely to control insects in agricultural practices and in health programs globally [5–7]. Sulfosulfuron, a new sulfonylurea herbicide is used to control weeds in wheat and other agricultural crops [8].

Pesticides are developed under strict vigilance of regulatory authorities which regulate the production of pesticides and set guidelines for their use so to put least hazardous impact on human health and on their surrounding environment. Besides these preventive measures, several health hazards can arise by their direct exposure during production and their contamination in food and water. Many environmental concern of pesticide has been already reported by researchers such as reducing biodiversity, nitrate leaching, decreased soil fertility, resistance in weed species towards common weedicides, increased cost of prevention and treatment endangering human health and acidification [9–10].

Several physical and chemical methods for degradation of the xenobiotic compounds are investigated but these methods are very costly and are not adequate enough [11–12]. Hence, the biological methods for degradation of xenobiotic compounds from the polluted sites by exploiting the microbial metabolism would be the best approach for the remediation of environment. The advantages of using this alternate remedial strategy (bioremediation) are, method is very cost effective, very economical, least hazardous, versatile and ecofriendly in nature [13–14]. Bacteria in nature degrade pesticides, and several toxic pollutants without any cost and do not cause secondary pollution. Consequently, researchers have conducted fine studies on biodegradation of toxic contaminants with clear understanding of their degradation mechanisms. Isolation of numerous bacteria and fungi efficient in degradation of pesticides into simple and non toxic forms have been done by various researchers [15–17].

Among the microbes used for biodegradation purposes, bacteria are most preferred as they could easily be developed into mutant strains with variety of biochemical pathways in adaptive environment [18–19]. Additionally, formulation of novel microbial consortia efficiently degrade xenobiotic compounds for their source of energy by using the co-metabolism process. The most promising and efficient technique for various biological purpose is the formation of proficient recombinant microbial strain which provides new opportunity to different biological fields [20].

Degradation of pesticides depends upon concentration, structure, solubility, soil types, soil moisture, temperature, pH, SOM (soil organic matter) and soil microbial biomass [12,21]. However, none of the studies have investigated degradation of different pesticides together through metagenomics tool to understand the mechanism behind *in situ* biodegradation and predict the biodegradation potential of microbes. Thus, there is a gap of knowledge regarding the functional genes and genomic potential underpinning degradation and community responses to contamination. Here we are trying to fill this knowledge gap by using advance technique like illumina sequencing of DNA isolated from pesticide contaminated agricultural soils and profiling of shifts in functional communities. Effect of pesticide stress on different category of proteins and their expression. On the basis of above reference the current study plans to the following objectives;

I. Isolation and characterization of Pesticide degrading bacteria
II. *In vitro* pesticide biodegradation in minimal medium
III. Identification of intermediate metabolites of biodegraded pesticide using GC-MS
IV. Estimation of pesticide degrading enzyme and their enzymatic kinetics
V. Proteomic analysis of pesticide degrading bacterial isolate in stress and normal condition
VI. Amplification of pesticide degrading gene
VII. Comparative metagenomic study of pesticide contaminated and uncontaminated soil

## Materials and Methods

### Chemicals and media used for experiment

Standard pesticides of highest purity namely cypermethrin, fipronil, imidacloprid and sulfosulfuron (Sigma-Aldrich 99% pure) were provided by Department of Chemistry, GBPUAT Pantnagar. Stock solutions (1mg/ml) of the pesticides were prepared by dissolving standard pesticide in acetonitrile (for non volatile compound or HPLC) or hexane (for volatile compounds or GC), then sterilized with the help of bacterial filter and stored in colored bottles at 4°C in refrigerator till use. Isolation and characterization of bacterial isolates was done using Nutrient Agar medium (NA) and Mineral salt medium (MSM).

### Isolation of Pesticide degrading bacteria through enrichment method

Pesticides polluted agricultural soil samples were collected from Gularbhoj, Udham Singh Nagar, Uttarakhand, India. Standard methods (Methods of Soil Microbiology and Biochemistry) were used to collect the soil samples [12]. For the collection of soil sample the handheld corer of stainless steel was used. During the sampling discarded the first two cores then third cores (5 cm), taken over an area of 100 m^2^ were pooled to form one composite sample. All the samples were packed in aluminium foil and sealed in sterilised plastic sampling bags to reduce the possibility of contamination. After proper labelling, samples were kept at −20°C until use. Isolation of pesticide degrading indigenous soil microorganisms was done by using enrichment technique followed by dilution method. For residual analysis of pesticides present in soil, the samples were prepared and performed according to Phartyal [22] and Negi et al. [23]. Extract of pesticides were analyzed by HPLC/GC. On the basis of the occurrence of residual pesticides in the soil sample, cypermethrin, fipronil, imidacloprid and sulfosulfuron were selected for the further study.

### Screening of pesticide degrading bacterial isolates

For the screening of cypermethrin, fipronil, imidacloprid and sulfosulfuron degrading bacteria, minimal salt medium was supplemented with different concentrations of the pesticide(s). Petri plates containing 20 ml of minimal agar supplemented with pesticide(s) (10 to 450 ppm from the stock solution (1mg/ml)) and left for solidification of medium. Plates were streaked with active bacterial cultures (isolated from the soil samples) and incubated for 72 h at 30±2°C. . The bacterial growth was observed after 2-4 days. On the basis of maximum tolerance level of the pesticides and growth in the minimal medium, bacterial isolates were selected, purified and maintained for further studies [12].

### Molecular characterization of pesticide degrading bacterial isolate

Selected bacterial isolate (2D) was characterized at morphological and molecular level. Genomic DNA was extracted from the bacterial isolate [24]. Amplification of 16S rDNA gene was done using universal primers (27f: 5′AGAGTTTGATCMTGGCTCAG3′ and 1492r: 5′TACGGYTACCTTGTTACGACTT-3′). After that the amplicon was processed for agarose gel electrophoresis and further sent for the 16S rDNA sequencing to Biotech Centre, South Campus, Delhi University. The contig of 16s rDNA gene was formed and check the percent similarity by comparing the sequences with NCBI gene database using BLAST programme. On the basis of maximum homology among the sequences the phylogeny of the organism was deduced with the help of MEGA 7.0 software [25].

### *In vitro* pesticide biodegradation in minimal medium

Experiment on biodegradation of cypermethrin, fipronil, imidacloprid and sulfosulfuron using selected bacterial isolates was performed in an *in vitro* study conducted under laboratory condition in liquid medium. Fifty ml MSM, supplemented with 20 ppm (1 ml from 1000 ppm pesticide stock) of pesticide individually was inoculated with 1 ml of active bacterial culture (OD=0.8) and incubated at 30°C. Further, t 2 ml of broth was withdrawn separately from all the flasks at different days interval (on 5, 10 and 15^th^ day) and centrifuged for 10 min at 10,000 rpm. After centrifugation, 1 ml of supernatant was added with 1 gm of sodium sulphate and 1 ml of hexane/acetonitrile. Two separate layers were formed in the separating funnel. Further the upper layer was collected and evaporated at room temperature with the help of evaporator. The dried pesticide in round bottom flask was next to mix properly with 2 ml of hexane/acetonitrile. After filtration extracted pesticide solution was collected and further analyzed with the help of GC/HPLC [12, 26].

### Identification of intermediate metabolites of biodegraded pesticide using GC-MS

On the basis of residual analysis of the pesticides in the medium using HPLC/ GC, percent biodegradation of cypermethrin was maximum among all the pesticides when treated with bacterial isolate (2D). Therefore the intermediate metabolites of cypermethrin biodegradation were analyzed with the help of GC-MS [27]. Degradation products of cypermethrin were extracted (50 ml of Mineral salt medium supplemented with 20 ppm of pesticide in 250ml flask containing) and analysed by GC-MS [12]. Uninoculated flask spiked with pesticide was used as control.

### Estimation of laccase enzyme in bacterial strain 2D

One ml of bacterial culture (12–14h old) inoculated into TYE broth (Tryptone yeast extract: tryptone, 2%; yeast extract, 2%; pH, 7.2) was incubated at 37 °C at 150 rpm for 5 days with (20 ppm) and without pesticide. Further, centrifugation at 6,000g was performed for 20 min at 4 °C. Before the sonication (at 20 MHz; five times, each for 45 s and 30 s gap between each step), 10 mM of PMSF (Phenylmethylsulfonyl fluoride) was added in 0.1 M phosphate buffer with pH 6.5 for washing cell pellets and inhibit protease activity in the supernatant. Again centrifugation at 14,000 g for 20 min at 4 °C was done to obtain cell extract which contains crude intracellular laccase enzyme. Enzyme activity can be measured at 465 nm using guaiacol as it forms reddish brown color in presence of laccase [28].

### Kinetics of laccase by Lineweaver-Burk model

Lineweaver-Burke plots were used to derive kinetic parameters of the laccase enzyme [12,29,30]. Equation 1 indicates Michaelis-Menton equation (where *Vmax* is maximum rate of reaction, *r* is substrate degradation in mM/min, *Km* is growth rate constant, *S* is initial substrate concentration).

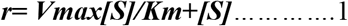

The inverse of Equation 1 is used in the Lineweaver-Burke plot:

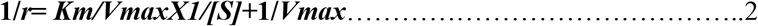

### Proteomic analysis of pesticide degrading bacterial isolate

To perform whole cell proteome analysis, bacterial strain was grown in minimal medium supplemented with 0.1% glucose in the presence/absence of the pesticide. Twenty ml of log phase bacterial culture, grown in minimal medium was used for the extraction of extracellular proteins. Bacterial culture was centrifuged at 12,000 rpm for 10 min at 4°C. Pellets were discarded and supernatant was used for further examination. Ammonium sulphate was added to precipitate the extracellular proteins. Suspension was kept at 4 °C overnight. Precipitated protein was concentrated by using dialysis. Concentration of protein was estimated [31]. Lyophilized protein samples were analysed using 2D gel electrophoresis. *In silico,* analysis of protein spots separated on 2D gel was carried with Expasy (TagIdent database) based on their molecular weight and isoelectric point (pI) [32].

### Amplifcation of laccase gene

Presence of laccase, a major pesticide degrading gene was targeted in the bacterial isolate and checked by using the primers [33]. The primers used for laccase amplification were CulAF-5′ACMWCBGTYCAYTGGCAYGG3′ and Cu4R-5′TGCTCVAGBAKRTGGCAGTG-3′. Reaction mixture constitutes: 20 pm/μl Primers (Forward and Reverse), 10 mM dNTPs mix, 10x Assay buffer with MgCl_2_, 3.0 U/μl TaqDNA polymerase, 10 mg/ml BSA 3 and 50ng/μl Template DNA. Reaction conditions maintained were: Initial denaturation at 94 °C for 3min followed by denaturation at 94 °C for30 sec, annealing at 50 °C for 30 sec, and from second phase repeat of 35 cycles, followed by extension at 72 °C for 1min and final extension at 72 °C for 5 min.

### Metagenomics of pesticide contaminated soil

#### Extraction of soil DNA

Collection of soil samples (pesticide contaminated and uncontaminated sites) was done from two different agricultural fields of Gularbhoj, Uttarakhand. Uncontaminated soil sample was taken as a control. DNA extraction from both samples was performed using HiPurA™ soil DNA Purification Kit. Soil (500g), from each sample was processed for DNA extraction. After quantification, purity of extracted DNA was tested in a NanoDrop spectrophotometer at 260 and 280 nm and by electrophoresis using 1% agarose. DNA concentration of the sample was 50 ng/L.

The microbiota of the soil samples was examined by targetingV3-V4 region of 16S rRNA through Illumina Miseq platform. Paired end reads obtained were further processed, checked (for score distribution, base quality, average base content and GC distribution) and merged using FLASH program [34]. Multiple filters were applied to generate high quality reads (~350 to ~ 450bp). UCHIME algorithm was used to detect and remove chimera. Entire downstream analysis was done using QIIME (Version 1.9.1) program [35]. All pre-processed reads obtained were pooled and clustered into OTUs (Operational Taxonomic Units) using Uclust program at 97% similarity. Representative sequences for each OTU were selected by aligning the sequences against Green gene data set via PyNAST [36]. Taxonomic classification was done by RDP classifier against SILVA 16S RNA genes database. Based on reads and OTU distribution of phyla and genera for each sample the reads were categorized. Alpha diversity was analyzed via three indices:chao1, shannon and observed-species matrix. All the indices were calculated using QIIME (Version 1.9.1).

### qRTPCR analysis

qPCR was carried out of 16S rDNA isolated from soilin iCycleriQ™ Multicolor instrument (Bio-Rad Lab, Hercules, CA, USA) using SYBR green dye [37]. A set of universal primers used were: (primer 1-5′ CCTACGGGAGGCAGCAG 3′and 2-5’ ATTACCGCGGCTGCTGG 3’).

## Results

Four pesticide utilizing bacterial isolates were recovered from pesticide contaminated agricultural fields using plate assay. Three bacterial isolates (2A, 2B and 2C) were able to grow at 300 ppm of the selected pesticides but only bacterial isolate (2D) was able to grow upto 450 ppm of the pesticides. Performance of 2D bacteria under laboratory condition (grew on cypermethrin, fipronil, imidacloprid and sulfosulfuron at 450 ppm) enabled us to investigate the biodegradation potential of the bacteria.

### *In vitro* biodegradation of pesticides in minimal medium

Gas chromatography (GC) results of standard of imidacloprid, fipronil and sulfosulfuron revealed the presence of major single peak of each pesticide at 4.3, 4.9, and 6.3 min (retention time) respectively under optimized conditions of the instrument. On the other hand cypermethrin showed four independent major peaks during a retention time of 16-17.65 **(Fig S1 a,b&c)**. These major peaks indicated at different retention time includes at 16.13 min indicate cis α, 17.00 min indicate cis β, 17.40 min indicate trans α and 17.62 min indicate trans β isomers of cypermethrin, **(Fig S1 d).** Isolate 2D showed maximum degradation of cypermethrin (94%) fo llowed by imidacloprid (91%), fipronil (89%) and sulfosulfuron (86%) after 15 days of incubation in minimal medium (**Fig 1)**.

**Fig 1.**
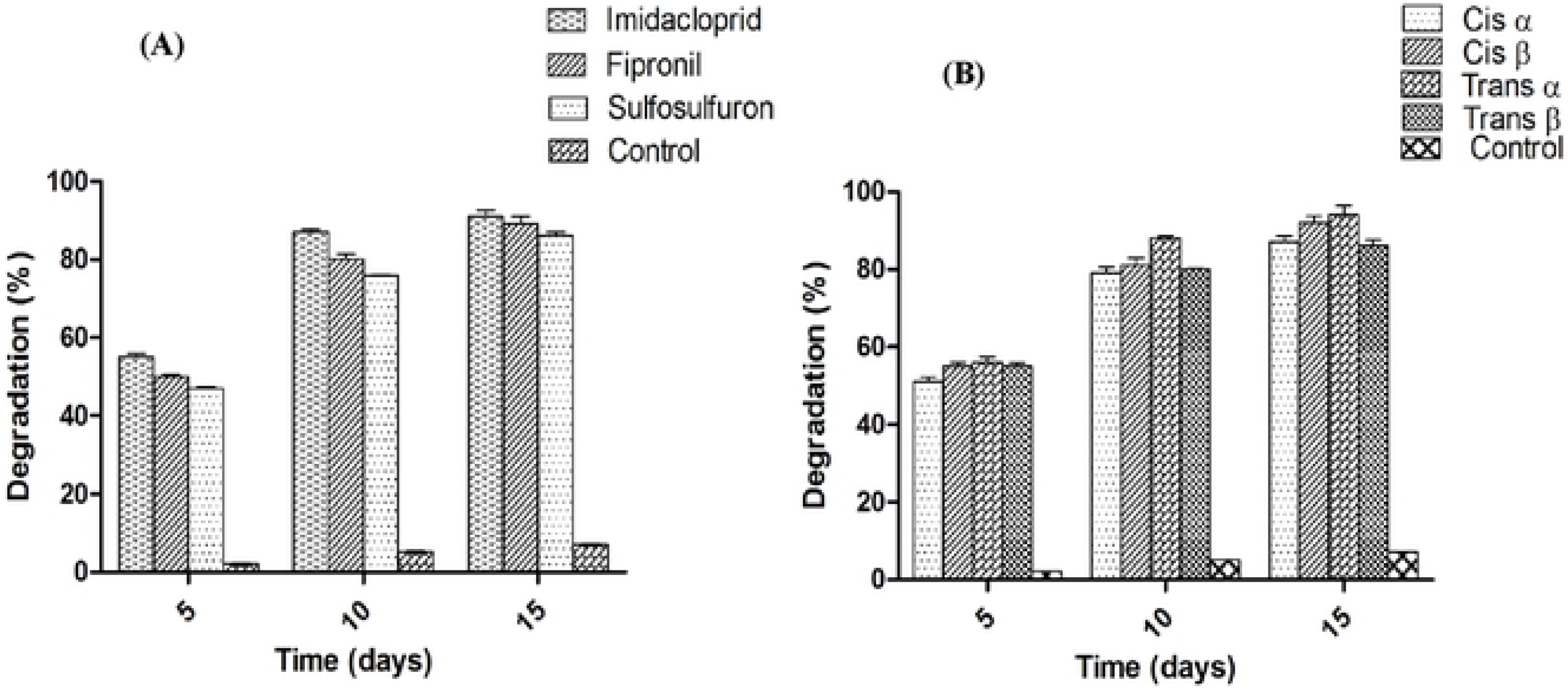
Biodegradation of imidacloprid, fipronil and sulfosulfuron and cypermethrin in minimal medium by 2D bacterial isolate. The graph denotes % degradation (Y axis) verses times in days (X-axis), where D denotes the days and numbers 5, 10 & 15 represent the withdrawl of sample(s) at different time interval. (A) represents the biodegradation of imidacloprid, fipronil, sulfosulfuron while (B) represents the biodegradation of different isomers of cypermethrin.

The intermediate metabolites of the cypermethrin were analyzed. Retention timings of different biodegraded products of cypermethrin under the treatment of bacterial isolate 2D were: 4.005, 25.018, 26.311, 28.286, 29.024 and 29.025 min which corresponded to 3-penten-1-ol and 1-[(1-oxo-2-propenyl) oxy]-2,5-pyrrolidin, 3-buten-2-one, 4,4-bis (dimethylamino) 4-quinolinol,1-ethyl-4-ethynyldecahydro, cyclobutane and pyrrolidine and chloroacetic acid& formic acid & decyl ester, 2h-pyrido[1,2-f][1,6] diazacyclooctadecin, acetic acid, n-[4-cyclooctylaminobutyl] aziridine & chloroacetic acid and octahydropyrrolo [1,2-a] pyrazine respectively **(Fig S2).**

### Characterization of the Bacterial Isolate

Phylogenetic analysis of amplified 16S rDNA gene sequences revealed 99% homology with *Bacillus cereus* of bacterial strain 2D. Gene sequences were submitted to GenBank and the organism was provided with an accession numbers (Accession No. MH341691) (**Fig S3 &S4)**.

### Enzyme kinetics

Laccase enzyme activity of test bacterial isolate was examined quantitatively and qualitatively for its possible role in degradation of cypermethrin. Biodegradation of pesticides is an enzyme-catalyzed transformation of organic compounds into their simpler products. Specificity of laccase enzyme make it crucial for degradation of huge variety of toxic environmental pollutants.

Michaelis Menten equation was solved in best way by plotting the data as 1/r vs 1/S to get a straight-line equation [29,38–39] **(Fig 2).** In the presence and absence of cypermethrin, bacterial isolate (2D) produced 163 and 29 μg/μL laccase respectively. Significant difference in *Km* values was observed with and without pesticide for laccase enzyme. *Km* values for laccase in 2D under stress and normal condition were 10.57 and 14.33 respectively.

**Fig 2.**
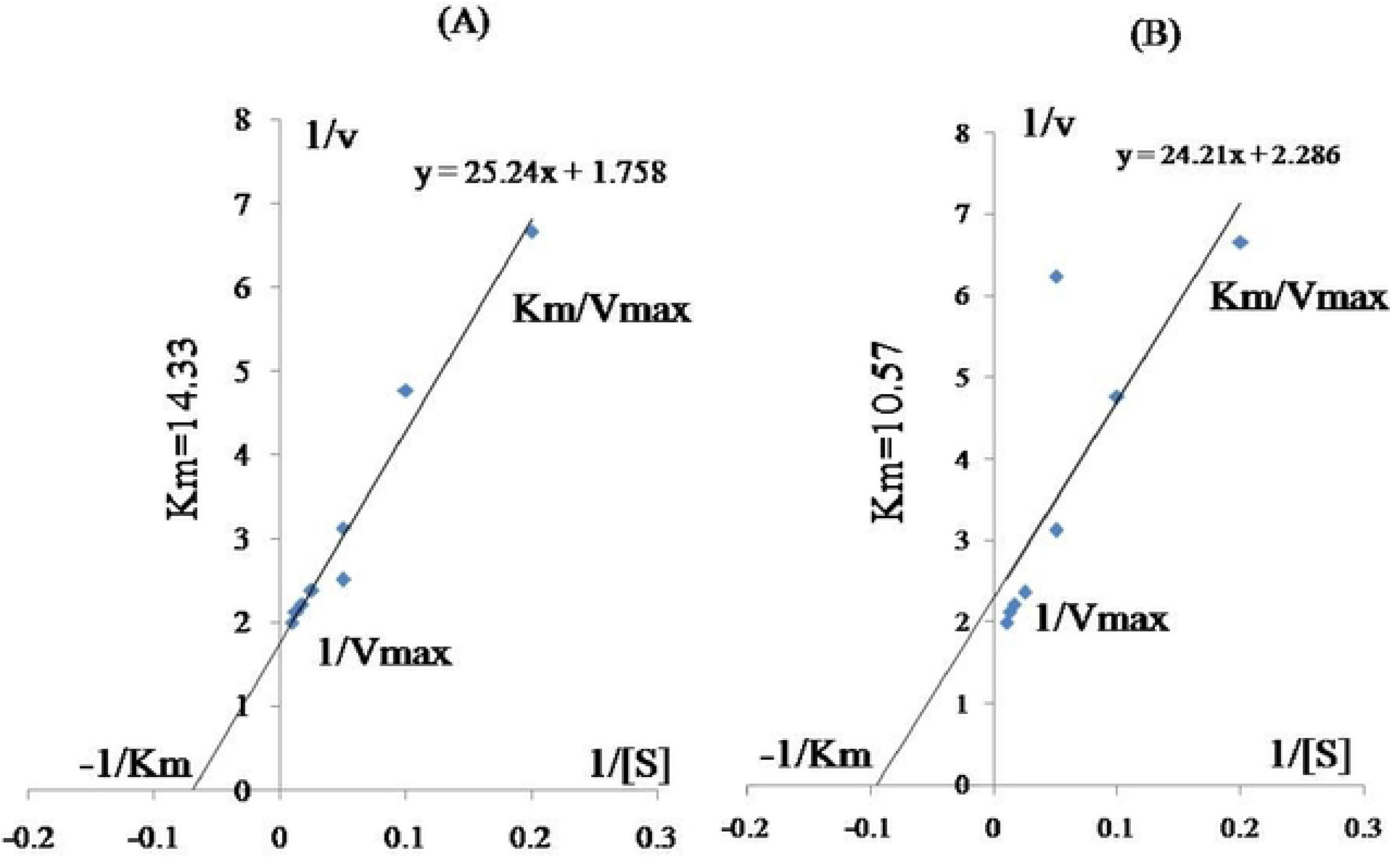
Lineweaver-Burke plot of laccase in bacterial isolates 2D (plot A and B). Plot A show laccase activity under normal condition while plot B shows laccase activity in minimal medium with pesticide.

### Laccase gene Amplification

Laccase amplification was found positive in test bacterial isolate and the size of amplicon for laccase was approximately 1200 bp **(Fig 3)**.

**Fig 3.**
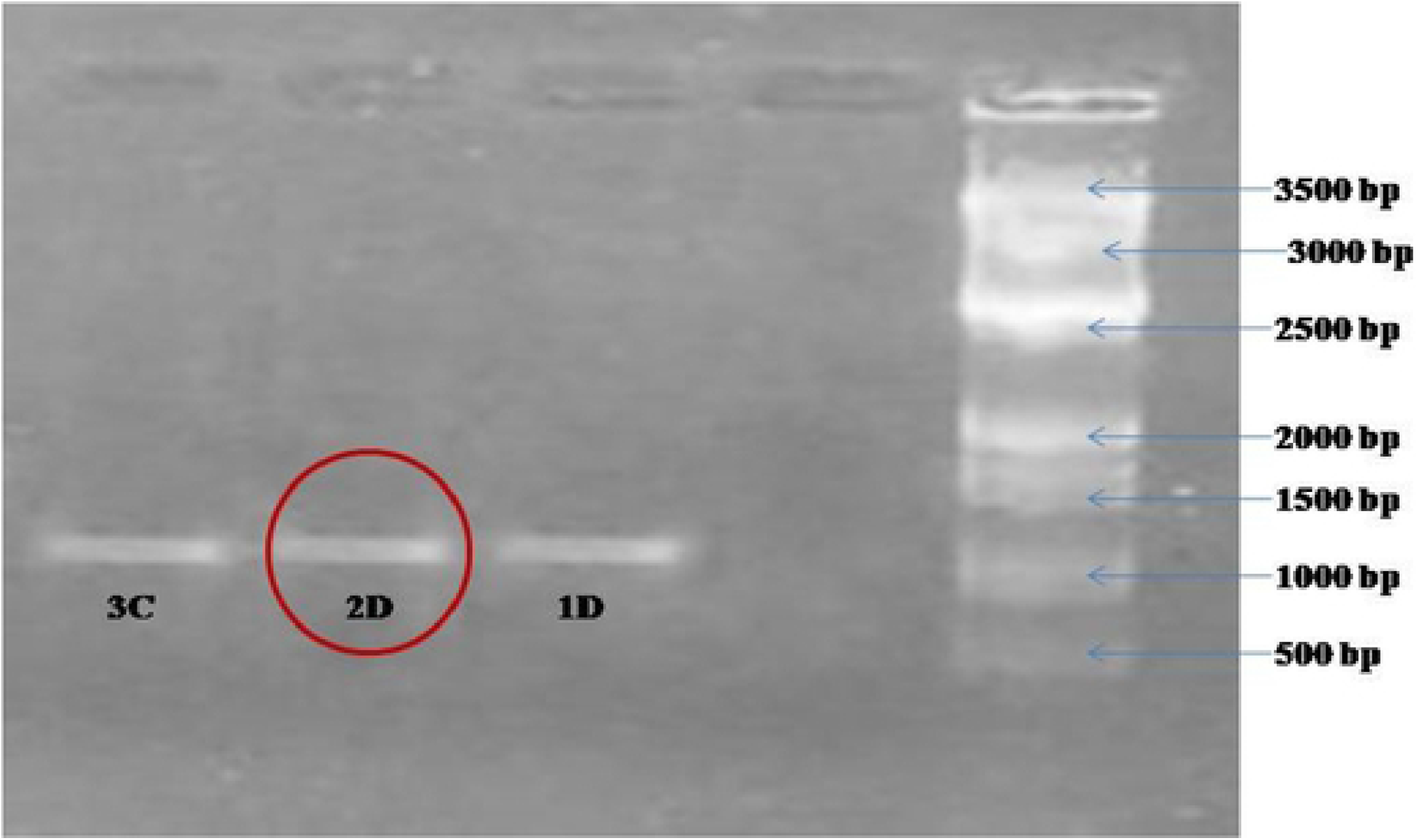
Amplification of Laccase gene from bacterial isolate 2D

### Comparative analysis of the pesticide induced proteins in 2D under stress and normal conditions

SDS PAGE was performed to analyse the expression of pesticide induced proteins in bacterial culture (2D) in the presence of cypermethrin (50 ppm). Size of different proteins bands were obtained ranged from 10 to 202 KD (**Fig S5**). Ten different protein bands were observed under stress condition as compared to normal condition. Further two-dimensional electrophoresis (2D electrophoresis) of the proteins was carried out in the absence and presence of pesticides **(Fig S6 & S7)** and major proteins were categorized in five classes namely: uncharacterized proteins, stress response proteins, catabolic proteins, energy production/ chemotaxis, gene regulation/ transcription and protein synthesis/ modification. Expression of protein by 2D was different under stress and normal conditions which could be differentiated on the basis of number of protein spots. Approximately 50 to 60 protein spots were found in the gel. We have characterized only 50 protein spots from each gel which were related to our experiment. Bacterial strain (2D) expressed 14% proteins for stress response, 32% proteins involved in protein synthesis and modification, 14% in gene regulation, 20% in energy production, 18% catabolic/pesticide degrading proteins and 2% were uncharacterized proteins under normal condition. Under stress, percent expression of protein was different and the order was: 30% stress response proteins, 22% of proteins involved in protein synthesis and modification, 12% in gene regulation, 14% in energy production, 22% catabolic/pesticide degrading protein and 2% uncharacterized proteins **(Fig 4).**

**Fig 4.**
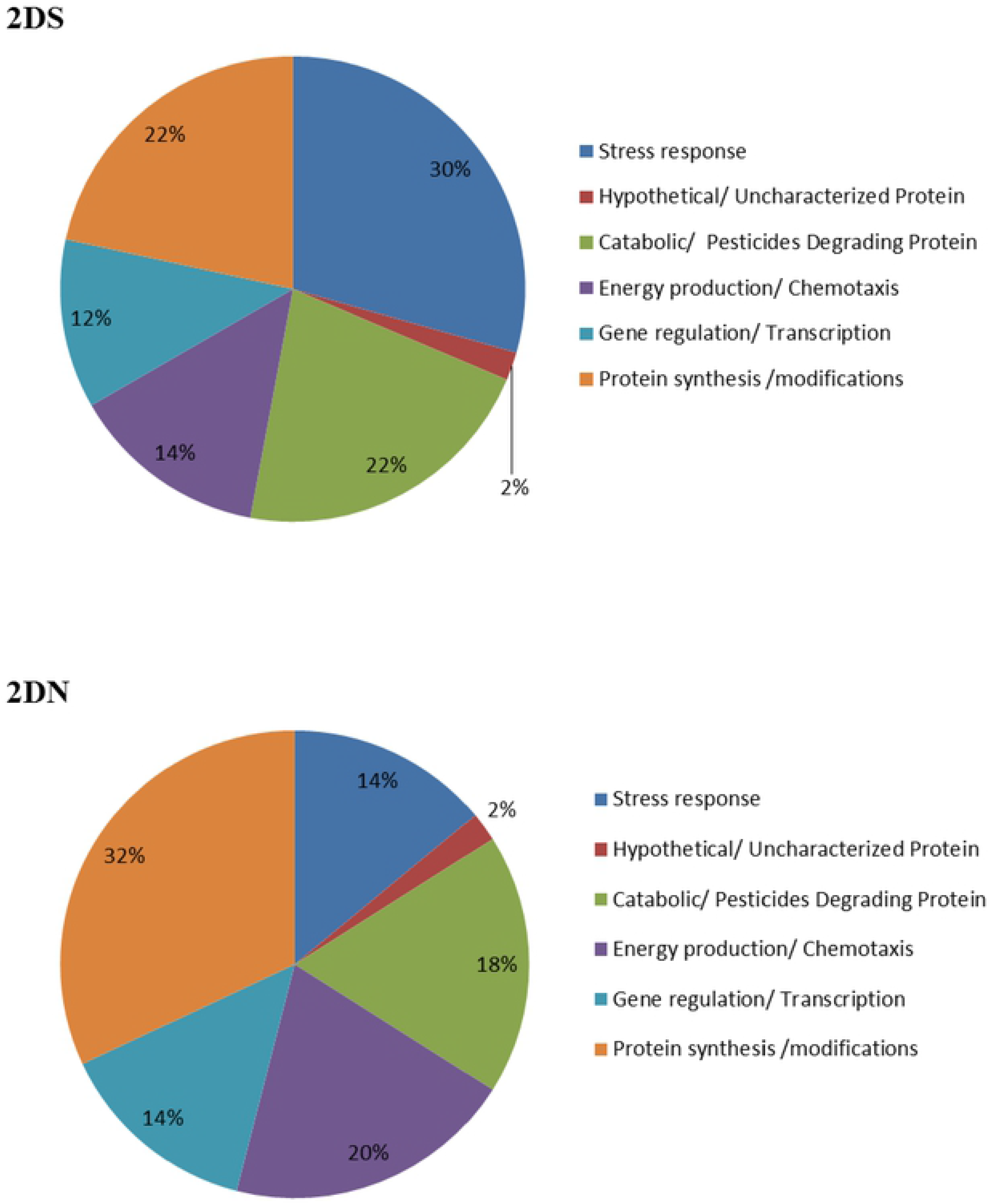
Expressed protein in normal (2DN) and pesticide stress condition (2DS) for bacterial strain 2D

### Real time PCR (qRTPCR) analysis

Maximum copy number of 16S rDNA per gram of soil sample was 1.96×10^8^ in pesticide contaminated soil while uncontaminated soil had 5.25×10^8^ copy number.

### Comparative analysis of microbial diversity in pesticide contaminated and uncontaminated soil samples

High-throughput Metagenomic approach was used to study the potential functional microbiome and composition of taxonomic community in pesticide contaminated soils.

### Statistics of Classification

Total reads in 2G (pesticide contaminated) and in 2GC (uncontaminated soil sample of Gularbhoj) were 562, 416 and 873, 083 respectively. A sum of 92.72% and 92.68% in 562, 416 reads (2G) and 873,083reads (2GC) respectively was classified at phylum level. The reads were further characterized at genus level and found to have 716 and 725 bacterial genera for 2G and 2GC soil samples respectively.

Community of abundantly present top 8 genera in both the soil samples was analysed. Unclassified category constitutes major genera in 2G (20.35%) and in 2GC (18.52%) soil samples. Second dominant genera in 2G soil were *Clostridium* (8.30%) followed by *Nocardioides* (3.41%) *Bellilinea* (3.14%), *Anaerolinea* (2.75%), *Longilinea* (2.48%), *Caldilinea* (2.33%) and *Phycicoccus* (2.21%). *Candidatus Koribacter* (5.89%) was second abundant genera in 2GC soil sample followed by *Bacillus* (5.06%), *Candidatus Solibacter* (5.00%), *Clostridium* (3.74%), *Conexibacter* (1.94%), *Streptomyces* (1.90%) and *Edaphobacter* (1.87%) **(Fig 5).**

**Fig 5.**
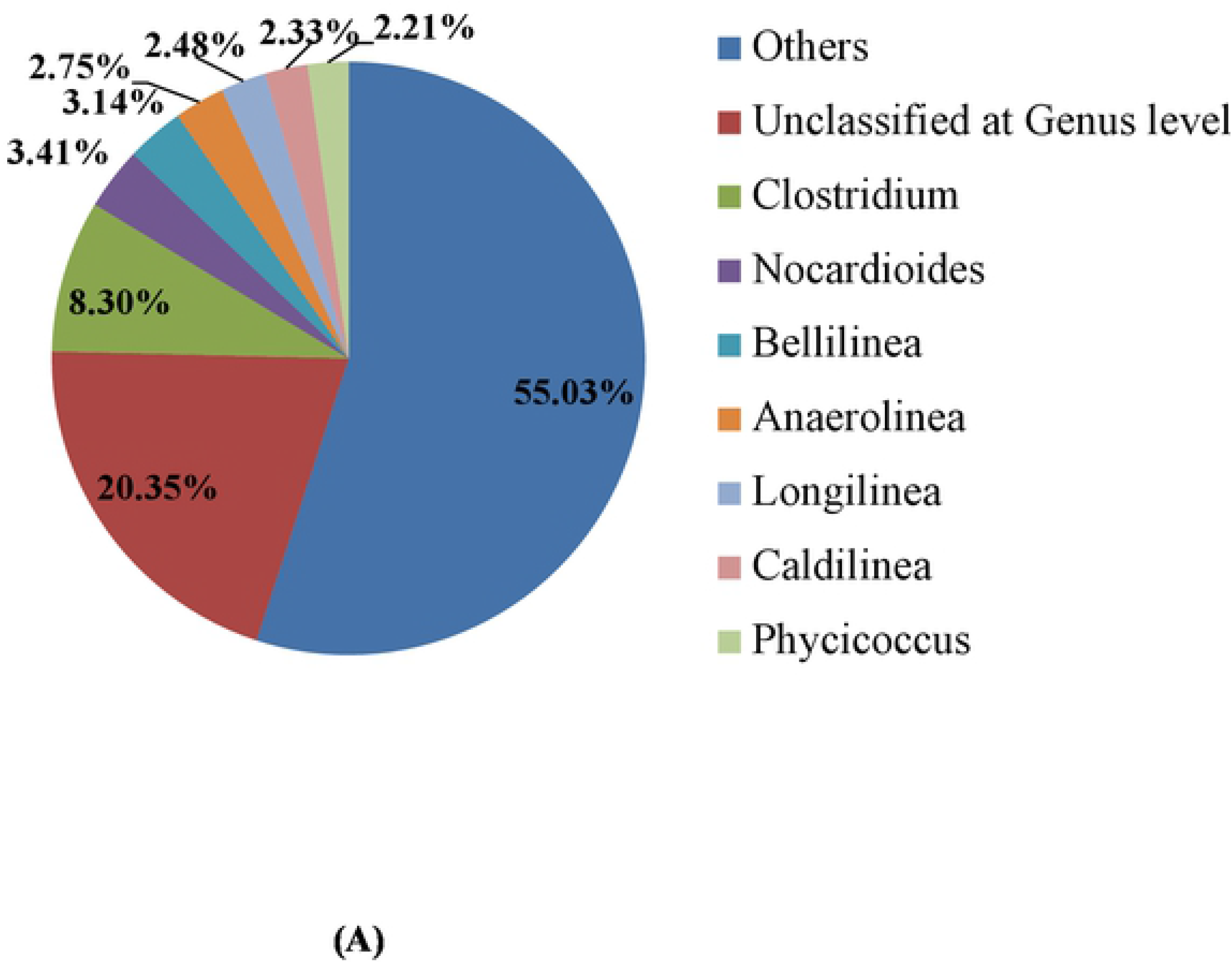
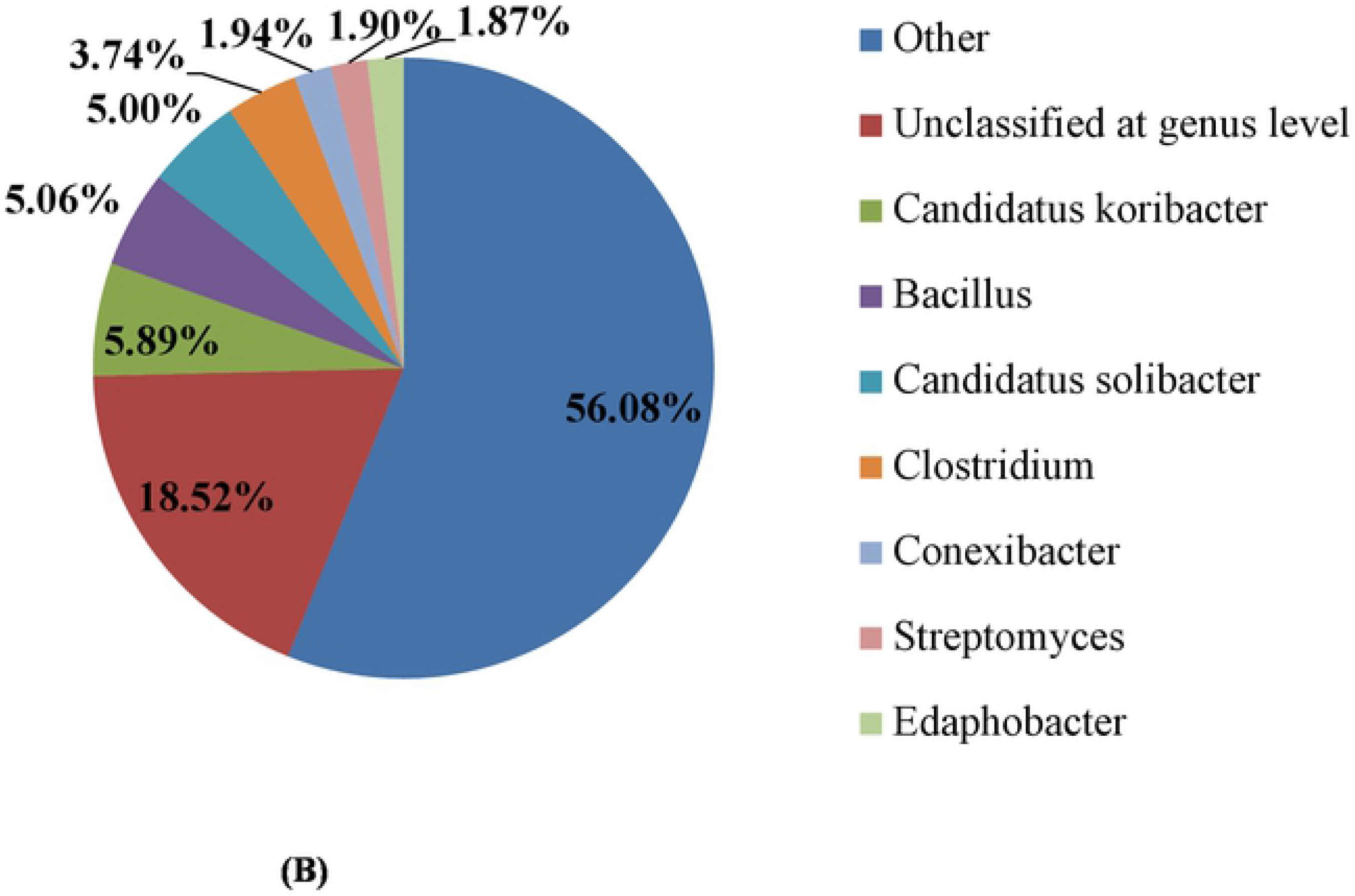
Pie chart represents the comparative analysis of 2G and 2GC soil samples at Genus level. [A] represents the 2G soil sample and [B] represents the 2GC soil sample.

Comparative analysis at Phylum level of both the soils revealed the presence of phyla viz. Firmicutes (*Clostridium*), Actinobacteria (*Nocardioides, Phycicoccus*) and Chloroflexi (*Bellilinea, Anaerolinea, Longilinea, Caldilinea*). While in 2GC soil, dominant Phyla were Acidobacteria (*Candida tuskoribacter, Candida tussolibacter*), Firmicutes (*Bacillus, Clostridium*), Actinobacteria (*Conexibacter, Streptomyces*) and Proteobacteria (*Edaphobacter*). Phyla namely Proteobacteria, Actinobacteria, Firmicutes, Bacteroidetes and Planctomycetes were invariably found in both the soil samples. Two unique Phyla: Chloroflexi and Nitrospira were present in 2G soil sample. Except *Clostridium,* overall composition of microbial communities was different at genus level in both the soil samples **(Fig S8).**

### Species diversity/ Alpha diversity

Relative abundance of top 25 classified OTUs was studied at genus level. Value of Shannon species diversity index was high (3.198) in pesticide contaminated (2G) soil. A sum of 1627 species were identified genotypically as compared to control (uncontaminated soil 2GC) where the Shannon species diversity index was 2.739 and 1,850 species were identified genotypically (**Fig 6)**.

**Fig 6.**
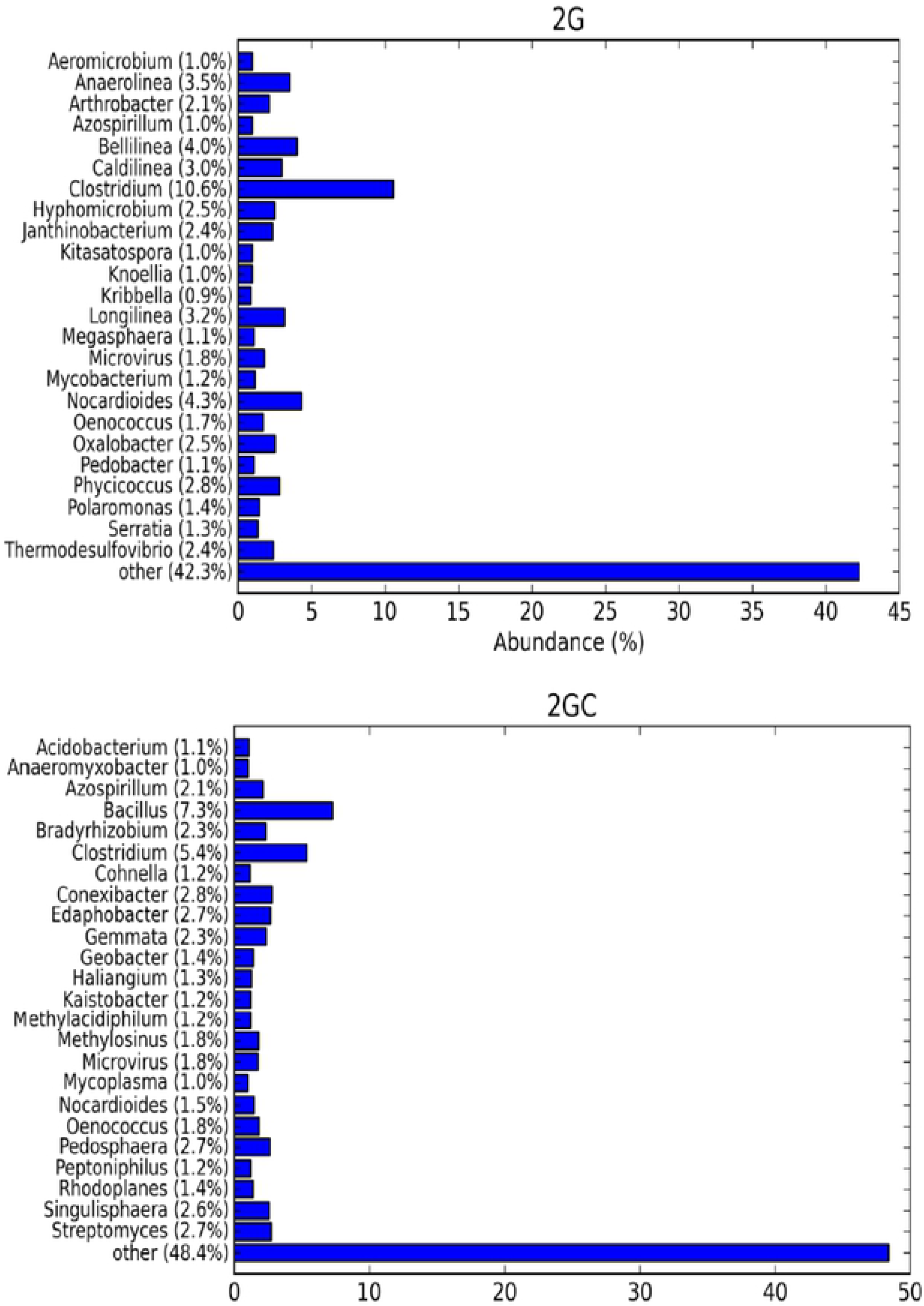
The bar graph show alpha diversity, the relative abundance of the top 25 classification OTUs results for genus taxonomic level of 2G and 2GC soil samples.

### Hierarchal clustering

The hierarchial clustering heat map displays OTUs count per sample. Higher the relative abundance of an OTU in a sample, the more intense the colour at the corresponding position in the heat map **(Fig 7).**

**Fig 7.**
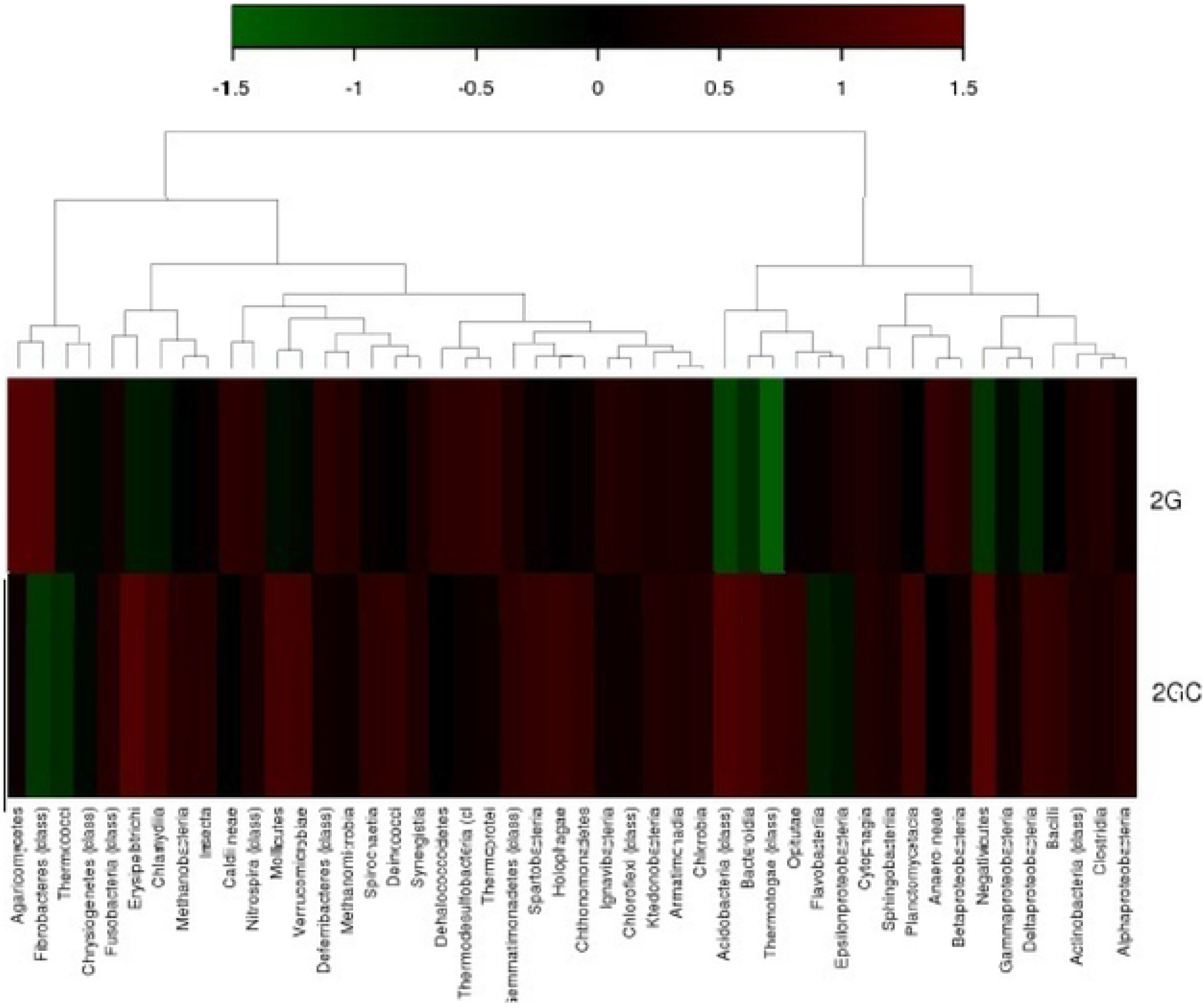
Hierarchial clustering represents total number of OTUs based on number of hits and % hits to the greengene database. Heatmap legends show the abundance of class in each sample. Deep brown (+1) to dark green (−1) show more abundance to less abundance distribution of OTUs.

The population size for *Clostridium* was larger in 2G but in 2GC soil sample the population size of *Koribacter* was large. Bacteria categorized as unclassified at genus level were dominantly present in both the soils but their percentage of abundance decreased in 2GC soil sample.

## Discussion

In the present study, interactions with the local farmers, survey of the nearby areas and the literature citations led to the findings that imidacloprid, cypermethrin, fipronil and sulfosulfuron are some common pesticides which are being used by the farmers in different cropping systems [40]. Presence of imidacloprid (0.42μg/g), cypermethrin (0.35μg/g) and fipronil (0.26 μg/g) and sulfosulfuron (0.18 μg/g) was found in soil samples. Degree of contamination in soil varies from less polluted (less then ≤ 0.5 mg/kg) to moderately polluted (in between 0.5 to 1 mg/kg) to heavily polluted (more then≥1 mg/kg) based on environmental quality standards [41]. Hence four pesticides (cypermethrin, fipronil, imidacloprid and sulfosulfuron) were selected in the present study for microbial biodegradation. The similar residual analysis of pesticides (carbendazim, endosulfan and imidacloprid) from the agricultural soil was done by the several researchers [22–23]. Our pesticide utilizing/ tolerant bacterial isolates were recovered from pesticide polluted agricultural soil using plate assay.

Performance of 2D bacteria under laboratory condition (grew on cypermethrin, fipronil, imidacloprid and sulfosulfuron at 450 ppm) enabled us to investigate the biodegradation potential of the bacteria. Cypermethrin, endosulfan, imidacloprid, fipronil and sulfosulfuron degrading bacterial and fungal strains were also isolated from the rhizospheric fields by several authors [12, 42–44].

Bacterial strain (2D) tolerated 450 ppm of all the four pesticides and degraded upto 20 ppm of the same in MSM medium. This indicates ability of bacterial isolate for pesticide degradation in different environments, as they can survive and degrade the toxic compounds on exposure of their higher concentrations. Recently similar study has been done in this field and found that *Bacillus spp*. tolerated carbendazim (150 ppm), *Microbacterium* tolerated 172 ppm of endosulfan *Pseudomonas* spp. tolerated 150 ppm of imidacloprid and *Pseudomonas* tolerated 30 ppm of lindane [23,45]. Therefore the bacterial isolate used in present study showed more tolerance for the pesticide(s) than those used by Nawab et al. and Negi et al. [23,45]. Bacterial strain 2D could degrade all the tested pesticides at higher concentration. Changes of regular expression of some enzyme(s) involved in degradation of imidacloprid, fipronil, cypermethrin and sulfosulfuron at increased concentrations by 2D cannot be ignored. On the basis of our experiment on tolerance for pesticides by 2D, we presume that metabolic activity of our strain was not subjected to catabolic repression at higher concentration of the pesticides. *Bacillus cereus* (strain 2D) potential of degradation and tolerance to pesticides (higher concentration) proved it as a suitable strain for remediation of polluted sites.

Isolated bacterial strain *Bacillus cereus* 2D showed higher range of pesticide degradation potential (94-86%) within shorter period of time (15 days). Pankaj et al. and Bhatt et al. [46,44] have reported biodegradation of cypermethrin using *Bacillus* spp. and found 85% degradation after 15 days under similar conditions. *Bacillus subtilis* BSF01 has been also reported for their schematic growth linked biodegradation pathway of β-cypermethrin [47]. Microorganisms used in biodegradation need an acclimatization period to induce synthesis of enzymes for biodegradation that may account for prolonged lag phase, which was observed at higher concentrations of cypermethrin, fipronil and sulfosulfuron [48]. *Bacillus* sp. FA3 degrades fipronil (76%) within 15 days, isolated from agricultural field and effective for its removal from water and soil environment [39]. Hence our bacterial strain *Bacillus cereus* 2D is more efficient than the previously reported bacteria for the removal of mixture of pesticide from the natural environment [39].

Toxicity of metabolites produced during cypermethrin degradation was checked by inoculating it (fifteen days old broth with metabolites) with fresh bacterial culture. Results proved these metabolites as nontoxic due to no effects on bacterial growth even after complete breakdown of pesticide. In the biodegradation experiment using 2D, a common intermediate of cypermethrin i.e., 3-Phenoxy benzoic acid was absent. It can be proposed that 3-PBA may be converted into smaller compounds like cyclobutane, chloroacetic acid and acetic acid. After molecular characterization of bacterial isolate 2D, it showed maximum similarity (99%) with *Bacillus cereus*. Several researchers displayed degradation abilities of *Bacillus* species of pesticides (such as profenos and cypermethrin) and other xenobiotic compounds [12,47,49–50].

Supplementation of pesticide with *Bacillus cereus* 2D, growing in minimal medium, showed reduced *Km* values as compared to higher *Km* values obtained under normal conditions, which indicates laccase production induced by addition of pesticide. This confirms competitive inhibition as the reason behind enzyme production. Laccase shows conformational changes, efficient activity and lower *Km* values in presence of pesticide. Increase in laccase activity and decreased km value under stress condition demonstrated that enzyme production increases under stressed conditions as compared to the normal condition. Production of laccase is high which helps bacterial isolates to overcome from the pesticide stress and make them comfortable to breakdown and utilize pesticides as carbon and energy source. Michaelis-Menten equation defines *Km* as the total substrate present at half of its maximum velocity (Vmax)[51]. Results indicate that to overcome the pesticide stress, the bacterial strain induces laccase enzyme. Oxidation of various inorganic and organic compounds (both phenolic and non-phenolic substrates) was catalysed by laccase *via* reduction of molecular oxygen to water [52]. Several biotechnological applications of laccase includes plastic degradation, xenobiotics bioremediation, bioleaching, decolourization of textile dyes, biosensors, lignin degradation, food industry, and biofuels production etc. [53–54]. Gangola and co-workers [12], reported efficient detoxification and degradation of cypermethrin by soil inhabitant *Bacillus subtilis* 1D with esterase and laccase activity. Laccase concentration in presence of cypermethrin was 62 μg/μL with *Km* value 61.57 M and in its absence was 42 μg/ μL with *Km* value 83 M after 15 days [12]. *Pseudomonas putida* MTCC 7525 was isolated from the soil sample containing sawdust and dairy effluent showed optimum laccase production (94.10 U/ml) at pH 8 and incubation at 30 °C with 1mM sodium nitrate as N source and 10% skim milk [55].

Presence of laccase gene was found in *Bacillus cereus* 2D and the size of amplicon for laccase was approximately 1200 bp **(Fig. 3).** Our results were similar to findings of Ausec et al. [33] as amplicon size of 1200 bp in *B. cereus* 2D was observed. By using primers it was confirmed that laccase encoding genes exists in bacterial genome and may get activated in stressed conditions in presence of pesticides). This gene is major regulatory gene and found responsible for the degradation of imidacloprid, fipronil, sulfosulfuron and cypermethrin. These pesticide have some common chemical bonds in their structures hence could be degraded by similar laccase. Presence of laccase gene has been already reported in *Azospirillum sp.* [56], *Pseudomonas syringae* [57] and *B. subtilis* [12]. Fungal laccase enzyme for degradation of liluron, chlorpyrifos and metribuzin was reported by Gouma, [58]. The release of extracellular laccase from *Bacillus cereus* allows degradation and decolorization of dye [59]. More studies were available on pesticide degradation by fungal laccase as compared to bacterial laccase. So, bacterial species with laccase activity makes them prominent candidate for degradation of variety of pollutants. The presence of laccase in *Bacillus cereus* (2D) is the first report in context of pesticide degradation.

Proteomic study revealed that under stress condition, percentage of stress responsive proteins and catabolic/pesticide degrading proteins was high, which make the organism more comfortable under such environment. There is direct involvement of proteins in pesticide degradation. Expression of such protein encoding genes have essential role in breakdown of toxic organic pollutant [60]. Proteomics explored degradation of PCB (polychlorinated bipheny) by microbes at molecular level [61]. Moreover, Pankaj and co-workers [62], reported expression of 250 proteins under stressed conditions (cypermethrin) and 223 proteins under normal conditions in *Bacillus spp.* These unique proteins were further classified into different groups based on their functionality, viz., translational proteins, stress proteins and catabolic enzymes etc. Up-regulation of proteins responsible for substrate transport and energy metabolism during pesticide degradation was reported in different microorganisms [7,46,63].

High- throughput metagenomic sequence analysis and qRTPCR were used to observe the impact of the pesticide on species richness and genetic diversity of microbes in the treated soil. 16S rDNA genes copy number in Gularbhoj soil (pesticide contaminated) suggested that the maximum number of pesticide degrading microbes are selectively enriched in the soil over other soil samples. Decrease in 16S rDNA genes copy number in Gularbhoj soil (pesticide contaminated) indicated negative effect of pesticide on microbial population. Investigation of gene expression and abundance of functional and taxonomic genes was done using Real time PCR [64]. Yale and co-workers [65] assessed and quantified expression of genes (trzN, atzB and atzA) in degradation of atrazine *via* Quantitative PCR (Q-PCR).

Decrease in reads of pesticide treated soil (2G) is supported by various findings which clearly denote the toxic nature of the pesticide. Decrease in read number in 2G soil sample may be related to excessive and repeated use of pesticides in the agriculture practices. Decrease in read number shows reduced microbial population. Pesticides had a negative effect on soil microorganisms on agar plates significantly [66–67]. Bacteria with potential to survive in pesticide treated soil were *Clostridium* sp., which is reported to degrade many pesticides like Alachlor, chlorpropham, DDT, Lindane etc. [68].

In 2G soil sample (pesticide contaminated), majority of genera were *Clostridium* (8.30%), *Nocardioides* (3.41%) *Bellilinea* (3.14%), *Anaerolinea* (2.75%), *Longilinea* (2.48%), *Caldilinea* (2.33%) and *Phycicoccus* (2.21%). *Nocardioides* was isolated from pesticide contaminated soil which utilized atrazine as a sole C and N source [69]. Presence of enzyme TrzN in *Nocardioides* sp. was responsible for biodegradation of pesticide chloro-s-trazina [70]. *Bellilinea* and *Longilinea* sp. are still not reported for pesticide biodegradation but *Bellilinea* sp. has shown catalytic breakdown of 2-methyl-naphthalene (a carcinogenic hydrocarbon) [67,71]. Thermophilic (*Bellilinea caldifistulae)* and mesophilic (*Longilinea arvoryzae)* strains were isolated from thermophilic digester sludge and rice paddy soil and found responsible for propionate-degradation. Role of *Anaerolineae* has been reported for wastewater treatment which had different recalcitrant hydrocarbons [72].

Shannon species diversity index was high (3.198) in pesticide contaminated (2G) soil as compared to uncontaminated soil (2GC) i.e., 2.739. Degradation effects of chlorpyrifos at variable concentrations on microbial diversity in soil has been studied *in vitro* which demonstrated similar variation in the diversity indices as found in biolog assay, showing chlorpyrifos inhibited microbial population in initial 2 weeks and later reached to same level as control [73].

Difference in microbial communities and population size indicates the repeated application of the pesticides in the agriculture field which forces the system to adopt the new population with replacement of the older one. The number of pesticide sensitive species decreased with time while pesticide tolerating species survived for longer. It is deduced from our results that pesticide is toxic for some species of the same phylum while other species utilize pesticide as a C and energy source and survived longer. Increase in population size of the Proteobacteria, Actinobacteria, Firmicutes and Chloroflexi showed that these are actively dividing bacteria in pesticide contaminated soil. Positive association of microbes in consortia may also intensify its abilities. We observed that communities with high metabolic potential for pesticide degradation were present in soils where pesticides retained for a longer time. Comparative investigation of two soil type suggested that abundance of high metabolic communities were present in rapidly degrading soil which indicates high functional capacity of the microbes in terms of nutrient cycling. We can say the microbial community present in 2GC was sensitive towards the pesticide while microbial community in 2G soil was resistant to pesticides.

Dominance of these genera (*Clostridium*, *Nocardioides*, *Bellilinea*, *Anaerolinea*, *Longilinea* and *Phycicoccus* in 2G soil sample) and presence of some unique genera in the soil samples shows that these microbes utilize pesticides as carbon and energy source for their growth. Pesticides were highly toxic for bacterial population hence some bacterial population was not present in treated samples but present in control sample. Higher richness of phyla Firmicute, Actinobacteria and Chlorofelxi in treated soil indicated their active participation in pesticide contaminated soil. Parks et al. and Jeffries et al. [74–75] used hierarchal map to study the relative abundance of microbial function in organophosphorus pesticide contaminated soil sample using UPGMA clustering. The hierarchial clustering heat map displays high abundance of class Clostridia, Actinobacteria, Betaproteobacteria, Cytophagia, Epsilonproteobacteria, Chloroflexi, Ignavibacteria, Chthonomonadeles, Holophagae, Chloroflexi, Gemmatimonadetes, Dehalococcoidetes, Caldineae, Thermodesulfobacteria and Thermoprotei in pesticide treated soil sample (2G).

## Conclusion

Present study recommends the application of indigenous microorganisms in biodegradation of common agriculture pesticides. Because in stress conditions bacteria could change their genetic profile and easily induce mutant strains, which can adopt in different environmental condition by activating their vast range of biochemical metabolism diversity, hence need to indepth study in this regard. The use of proteomics tools for the purpose of environmental bioremediation gives detailed information of microbial cells protein and composition. Hence it offers a valuable approach to decipher the mechanisms involved in bioremediation at molecular level. Diversity, composition and metabolic potential of soil microbiome is of crucial importance in bioremediation. In addition soil microbial communities also regulate biogeochemical cycling. Hence, detailed analysis of microbial communities at functional and structural level will guide to monitor and assess the effect of pesticides on soil health and their biological status respectively. Therefore, based on the biodegradation potential and wide range of activity for different substrates, the *Bacillus cereus* 2D could be a solid recommendation for the biodegradation of pesticides. While the Metagenomic study of contaminated sample (soil or water) could unrevealed the type of microbial population and the interaction among them for the need of pesticide biodegradation in natural environment. For future prospects the activity of bacterial isolate could be check against the other xenobiotic compounds and the mechanism of gene regulation.

## Acknowledgements

Authors are acknowledging the Department of Microbiology and Department of Chemistry, G.B. Pant University of Agriculture and Technology, Pantnagar for providing all the laboratory facility during the study.

## Authors’ contributions

Saurabh Gangola: Planned and designed the experiments, participated in all the experiments and created the manuscript; Samiksha Joshi, Saurabh Kumar: Participated in all the experiments and edited the manuscript; Anita Sharma: Supplied the experimental instruments and laboratory facilities. All authors have read and approved the manuscript.

## Additional information

**Supplementary files**: Supplementary file is attached for the support of data. The Accession number of the isolated strain 2D is MH341691 provided from NCBI

## Notes

### Competing Interest Statement

The authors have declared no competing interest.

